# Enzymes enabling the biosynthesis of various C_20_ polyunsaturated fatty acids in a sea urchin *Hemicentrotus pulcherrimus*

**DOI:** 10.1101/2024.05.23.595617

**Authors:** Yingying Peng, Yutaka Haga, Naoki Kabeya

**Author notes:** Corresponding author Naoki Kabeya.

## Abstract

Sea urchins, integral to marine ecosystems and valued as a delicacy in Asia and Europe, contain physiologically important long-chain (>C_20_) polyunsaturated fatty acids (PUFA) in their gonads, including arachidonic acid (ARA, 20:4n-6), eicosapentaenoic acid (EPA, 20:5n-3), and unusual non-methylene interrupted fatty acids (NMI-FA) such as 20:2^Δ5,11^. Although these fatty acids may partially be derived from their diet, such as macroalgae, the present study on *Hemicentrotus pulcherrimus* has uncovered multiple genes encoding enzymes involved in LC-PUFA biosynthesis. Specifically, three fatty acid desaturases (FadsA, FadsC1 and FadsC2) and thirteen elongation of very-long chain fatty acids proteins (Elovl-like, Elovl1/7-like, Elovl2/5-like, Elovl4-like, Elovl8-like, Elovl6-like A to H) were identified in their genome and transcriptomes. Functional analysis showed that FadsA and FadsC2 function as a Δ5 desaturase and a Δ8 desaturase, respectively, enabling the conversion of 18:2n-6 and 18:3n-3 into ARA and EPA, respectively, along with Elovl, particularly Elovl6-like C. Elovl6-like C demonstrates elongase activity towards both C_18_ PUFA and monounsaturated fatty acids (MUFA). Consequently, FadsA and Elovl6-like C enable to synthesise several NMI-FA, including 20:2^Δ5,11^ and 20:3^Δ5,11,14^, from C_18_ precursors. This indicates that *H. pulcherrimus* can endogenously synthesise a wide variety of C_20_ PUFA and NMI-FA, highlighting active biosynthesis pathways within sea urchins.

## 1. Introduction

Long-chain (≥C_20_) polyunsaturated fatty acids (LC-PUFA) are important components of cell membranes in animals and play a crucial role in maintaining their structural and functional integrity (Yehuda et al., 2002). In particular, eicosapentaenoic acid (EPA, 20:5n-3) and docosahexaenoic acid (DHA, 22:6n-3) play beneficial roles in prevention of various human diseases, including cardiovascular disease, inflammatory diseases like rheumatoid arthritis, Crohn’s disease and ulcerative colitis, as well as types of cancer such as colorectal, breast and prostate cancers. They also ensure normal cognitive and visual function during early development (Brouwer et al., 2006). All deuterostomes including vertebrates and echinoderms are incapable of *de novo* PUFA biosynthesis from non-PUFA precursors due to the absence of specific enzymes (Kabeya et al., 2018), and thus require pre-formed C_18_ polyunsaturated fatty acids (PUFA) in their diets to synthesise LC-PUFA endogenously. In most eukaryotes, two types of enzymes, named front-end desaturases (Fads) and elongation of very-long chain fatty acids proteins (Elovl) are required to synthesise LC-PUFA from C_18_ PUFA (Supplementary Fig. 1). The former introduces a new unsaturation (C=C double bond) between the pre-existing one and the carboxy-terminus in the fatty acyl chain, and the latter are rate-limiting enzymes catalysing the initial condensation step of fatty acid elongation to add a C_2_ unit. Elovl can further be divided into two major subgroups, according to their sequences and overall substrate preferences on either saturated/monounsaturated fatty acids (S/MUFA) or PUFA, which have been determined in vertebrates (Hashimoto et al., 2008). Fads also exhibit different regioselectivities to add a double bond at a specific carbon in the fatty acyl chain. Indeed, their substrate specificities and complement of genes encoding those enzymes vary among different organisms, leading to remarkable diversity in the series of LC-PUFA that can be endogenously synthesised in specific animals. The ever-increasing availability of genomic and transcriptomic information in non-model organisms has also driven the elucidation of LC-PUFA biosynthetic capacity in a variety of animals, including invertebrate species (Monroig & Kabeya, 2018).

Echinoderms, a group of marine invertebrates, include extant species such as sea urchins (Echinoidea), sea cucumbers (Holothroidea), sea stars (Asteroidea), brittle stars (Ophiuroidea) and sea lilies (Crinoidea). In recent years, sea urchins have gained global attention as a premium commodity. Both male and female sea urchins are harvested for their gonads, typically referred to as “roe” in the fishery and catering markets (Carboni et al., 2013; Rahman et al., 2014). In countries across Asia, the Mediterranean and the Western Hemisphere, sea urchin roe is considered a prised delicacy, particularly valued in Japanese sushi (Rahman et al., 2014; Sun & Chiang, 2015). Fatty acid analysis has shown that commercially important sea urchins contain various PUFA, notably arachidonic acid (ARA, 20:4n-6) and EPA (Zhukova, 2022). Although sea urchins primarily consume macroalgae which contain low amounts of LC-PUFA such as ARA and EPA, it appears these LC-PUFA might be accumulated from their diets (Castilla-Gavilán et al., 2018; Miyashita et al., 2010). However, a previous study on the sea urchin *Paracentrotus lividus* has demonstrated that they possess three Fads (FadsA, FadsC1 and FadsC2), which suggest a capacity for endogenous LC-PUFA biosynthesis (Kabeya et al., 2017). Evidence from feeding trials has shown that certain sea urchins such as *P. lividus* and *Strongylocentrotus droebachiensis* were able to survive and grow normally on diets free of LC-PUFA, typically land vegetables (Fernandez & Boudouresque, 2000; Liyana-Pathirana et al., 2002; Vizzini et al., 2015). This raises questions about whether they possess a complete gene set required for the full biosynthetic pathway of LC-PUFA including the physiologically important ARA, EPA and DHA from their C_18_ PUFA precursors. In addition, functional analysis of FadsA from *P. lividus* revealed its capability to synthesise unusual PUFA named non-methylene interrupted fatty acids (NMI-FA) from C_20_ PUFA substrates by introducing a double bond at the Δ5 position in the carbon chain. These NMI-FA, such as 20:3^Δ5,11,14^ and 20:4^Δ5,11,14,17^ are indeed commonly detected in several Echinodermata species including sea urchins *Anthocidaris crassispina*, *Strongylocentrotus intermedius*, *Strongylocentrotus nudus*, *Hemicentrotus pulcherrimus* and *Pseudocentrotus depressus* (Takagi et al., 1986; Zhukova, 2022), a sea star *Asterias vulgaris* (Paradis & Ackman, 1977; Zhukova, 2022) and a brittle star *Ophiura sarsi* (Sato et al., 2001). Two other NMI-FA (20:2^Δ5,11^ and 20:2^Δ5,13^) are also prevalent in sea urchins, with NMI-FA levels in *S. droebachiensis*, *S. intermedius* and *P. depressus* comparable to EPA levels (Takagi et al., 1986; Kelly et al., 2008; Zhukova, 2022). The physiological roles of these NMI-FA remain unclear, but their relatively high levels suggest they may play significant roles in echinoderms.

In the present study, we comprehensively isolated a series of *fads*- and *elovl*-like sequences from the transcriptomic and genomic databases of the sea urchin *H. pulcherrimus*. Subsequently, sequence and phylogenetic analysis were conducted to determine the orthology of these genes with those found in vertebrates. Furthermore, the obtained sequences were heterologously expressed in yeast for the functional analysis to determine the biosynthetic capabilities of LC-PUFA in *H. pulcherrimus*.

## 2. Materials and Methods

### 2.1. Extraction of total RNA and cDNA synthesis

*H. pulcherrimus* (weight 11.6 g, female) used in the present study were randomly collected from intertidal areas in Tateyama Station, The Field Science Center, Tokyo University of Marine Science and Technology (34.9763, 139.7699, Tateyama, Chiba, Japan) on May 6, 2022. Subsequently, total RNA was extracted from their gonad and intestine using TRIzol (Thermo Fisher Scientific, Waltham, MA, USA) following the manufacturer’s recommendations. Complementary DNA (cDNA) was synthesized from 5 µg of the total RNA using the SuperScript IV Reverse Transcriptase Kit (Thermo Fisher Scientific) following the manufacturer’s recommendations.

### 2.2. In silico isolation of elovl- and fads-like sequences from H. pulcherrimus genome and transcriptome

The *elovl*- and *fads*-like sequences of *H. pulcherrimus* were comprehensively retrieved from their genome and/or transcriptome assemblies (NCBI WGS project no. BEXV01, TSA project no. IACU01) by *tblastn* using several known Elovl and Fads sequences, respectively, as queries. Three *fads*- and eight *elovl*-like sequences were identified by this search as possible enzyme genes involved in LC-PUFA biosynthesis. Based on those putative sequences, primer sets for the full-length open reading frame (ORF) cloning were designed with the corresponding restriction enzyme site to ligate amplified ORF into the yeast expression vector pYES2 (Thermo Fisher Scientific). All primer sequences used in the present study were listed in Supplementary Table 1.

### 2.3. Phylogenetic analyses of sea urchin Fads and Elovl

The *elovl*-like sequences from Echinodermata species were retrieved from NCBI nr and TSA databases using *blastp* and *tblastn*, respectively with several Elovl sequences from vertebrates as queries. The full-length amino acid (aa) sequences were selected from the sequences retrieved from nr database. Regarding the transcriptomic sequences from TSA database, the sequences containing full-length ORF were selected and translated into deduced aa sequences. After the selection, we divided all Elovl sequences into two groups, namely S/MUFA subfamily (S/MUFA Elovl) and PUFA subfamily (PUFA Elovl) according to Hashimoto et al. (2008); S/MUFA Elovl and PUFA Elovl should contain H-W/T-X-H-H and Q/H-X-T/S-X-L-H-X-X-H-H, respectively (Supplementary Table 1). In order to determine orthologues of vertebrate Elovl, human and zebrafish Elovl aa sequences were added to the dataset for the phylogenetic analysis described below. An initial retrieval method of Fads sequences from Echinodermata species were the same with that used for Elovl sequences, although query sequences were several Fads sequences from a sea urchin *P. lividus* and vertebrates. The collected Fads-like sequences were then screened by using three histidine box sequences which are well-conserved among Fads (i.e., H-X-X-X-H, H-X-X-H-H and Q-X-X-H-H) (Supplementary Table 1). For the maximum likelihood phylogenetic inference, multiple sequence alignments (MSA) were created by using MAFFT v7.407 with *einsi* mode (Katoh et al., 2017). The resulting MSA were then trimmed by trimAl to remove any columns containing gaps which were >95% of the sequences (Capella-Gutierrez et al., 2009). After the best-fit aa substitution model was selected as JTT+I+G4+F for S/MUFA Elovl, LG+I+G4 for PUFA Elovl and Fads by ModelTest-NG (v0.1.6) (Darriba et al., 2019), the phylogenetic inferences were performed using RAxML-NG (v1.2.0) with *--all* option (ML search + bootstrapping) (Kozlov et al., 2019). The resulting ML trees were visualised using CLC Main Workbench v21 (Qiagen).

### 2.4. Functional characterisation in the yeast Saccharomyces cerevisiae

PCR for the full-ORF amplification were performed with PrimeSTAR Max DNA Polymerase (TaKaRa Bio Inc. Shiga, Japan) following the manufacturer’s recommendations. Briefly, a thermocycler condition was set to 98°C for 2 minutes as the initial denaturation, 35 cycles of 98°C for 10 seconds, 55°C for 5 seconds and 72°C for 15 seconds, and then 72°C for 2 minutes as the final extension. Subsequently, the amplified PCR products were digested using the corresponding restriction enzymes (Supplementary Table 1) and then ligated into similarly digested pYES2 using LigaFast Rapid Ligation System (Promega Corporation, Madison, WI, USA) following the manufacturer’s recommendations. The ligated product was transformed into *E. coli* using ECOS Competent *E. coli* DH5α (Nippon Gene Co., Ltd., Tokyo, Japan) following the manufacturer’s recommendations. After confirming sequence (Eurofins Genomics K.K, Tokyo, Japan), the successful plasmid DNA was transformed into the INV*Sc*1 yeast (Thermo Fisher Scientific) using *S.c.* EasyComp^TM^ Transformation Kit (Thermo Fisher Scientific) following the manufacturer’s recommendations. The yeast transformation and culture conditions were described previously (Kabeya et al., 2021). Each transgenic yeast was grown in the presence of one of the potential PUFA substrates for 48 h at 30°C. For Elovl, the exogenously added PUFA substrates were 18:2n-6, 18:3n-3, 18:3n-6, 18:4n-3, 20:4n-6, 20:5n-3, 22:4n-6 and 22:5n-3. We also grow the transgenic yeast without any of substrate PUFA, only when Elovl have shown the activity towards the yeast endogenous SFA and/or MUFA. For Fads, the PUFA substrates were 18:2n-6, 18:3n-3, 20:2n-6, 20:3n-3, 20:3n-6, 20:4n-3, 22:4n-6 and 22:5n-3. In addition to PUFA substrate, 20:1n-7 and 20:1n-9 were also used as the exogenously added substrates for Fads to test Δ5 desaturation activity to produce non-methylene interrupted fatty acids. The yeast transformed with empty pYES2 were also cultured as a negative control. The resulting yeast culture were collected and then lyophilised for the preparation of fatty acid methyl ester (FAME) which were prepared as previously described (Kabeya et al., 2021). All fatty acid substrates were purchased from Nu-Chek Prep, Inc. (Elysian, MN, USA), except 18:4n–3 from Matreya, LLC. (State College, PA, USA) and 20:1n–7, 20:1n–9 and 20:4n–3 from Larodan AB (Solna, Sweden).

### 2.5. Fatty acid analysis by a gas chromatography

The FAME prepared from the yeast were analysed by a gas chromatograph (Nexis GC-2030; Shimadzu Corporation, Kyoto, Japan) equipped with a flame ionisation detector (FID) and a capillary column (FAMEWAX; 30m x 0.25 mm, i.d. x 0.25 μm, Restek, Bellefonte, PA, USA). The injection port and FID temperatures were set to 250 °C and 280 °C, respectively. A constant velocity (40 cm/s) was delivered with hydrogen as a carrier gas. The column oven temperatures were programmed for an initial 50 °C for 1 min, elevating from 50 °C to 190 °C at a rate of 40 °C/min and then 190 °C to 240 °C at a rate of 4 °C/min with holding at a final temperature of 240 °C for 3 minutes. Each detected peak was identified by comparing the retention times of the samples with those obtained in commercial standards. The conversions (%) of each enzyme were calculated according to a formula: [all product areas/ (substrate area + all product areas) × 100].

### 2.6. Structural determination of non-methylene interrupted fatty acids by a gas chromatography mass spectrometry

To determine the structure of unusual NMI-FA detected in the yeast assay, we prepared 4,4-dimethyloxazoline (DMOX) derivatives from the FAME samples following the previously described method (Kabeya et al., 2017) with a slight modification. Briefly, the FAME samples were transferred into a glass ampoule and 2-amino-2methyl-1-proponol was added. After closing the ampoule, the mixtures were incubated over-night at 180°C. The resulting DMOX derivatives were extracted by diethyl ether-hexane (1:1, v/v), dried-up and then dissolved in hexane to be injected into a gas chromatograph mass-spectrometer (GCMS-QP2010; Shimadzu Corporation) equipped with a capillary column (SUPELCOWAX™ 10; 30 m x 0.32 mm i.d. x 0.25 μm, Merck KGaA, Darmstadt, Germany). The injection port and ion source temperatures were 250 °C and 200 °C, respectively. Helium was used as a carrier gas with a constant velocity (60 cm/s). The programme of column oven temperatures was set to 50 °C for 1 min, elevating from 50 °C to 180 °C at a rate of 40 °C/min and then 180 °C to 240 °C at a rate of 1/min.

## 3. Results

### 3.1. Sequence and maximum likelihood phylogenetic analysis of elovl-like and fads-like sequences isolated from H. pulcherrimus

Thirteen *elovl*-like sequences were successfully isolated from *H. pulcherrimus*; the eight and five out of thirteen were classified as the S/MUFA Elovl and PUFA Elovl, respectively, according to their amino acids sequences (Supplementary Table 1). Regarding S/MUFA Elovl, we named them as Elovl6-like A to H since Elovl6 is a commonly conserved S/MUFA Elovl among vertebrates (Fig. 1A). Four out of five PUFA Elovl from *H. pulcherrimus* formed relatively well-supported clades with each PUFA Elovl from vertebrate species in the ML phylogenetic tree and accordingly, we named them Elovl2/5-like, Elovl4-like, Elovl1/7-like and Elovl8-like (Fig. 1B). However, since one PUFA Elovl from *H. pulcherrimus* was not clustered together with any of vertebrate Elovl in the ML phylogenetic tree, we named it Elovl-like in the present study. Regarding *fads*, three full-length ORF sequences were successfully isolated from *H. pulcherrimus*. The ML phylogenetic analysis of echinoderm Fads demonstrated that each of *H. pulcherrimus* Fads was clustered together with either FadsA, FadsC1 and FadsC2 from *P. lividus*, and thus the corresponding names were given to each of *H. pulcherrimus* Fads (Fig. 2).

**Fig. 1.**
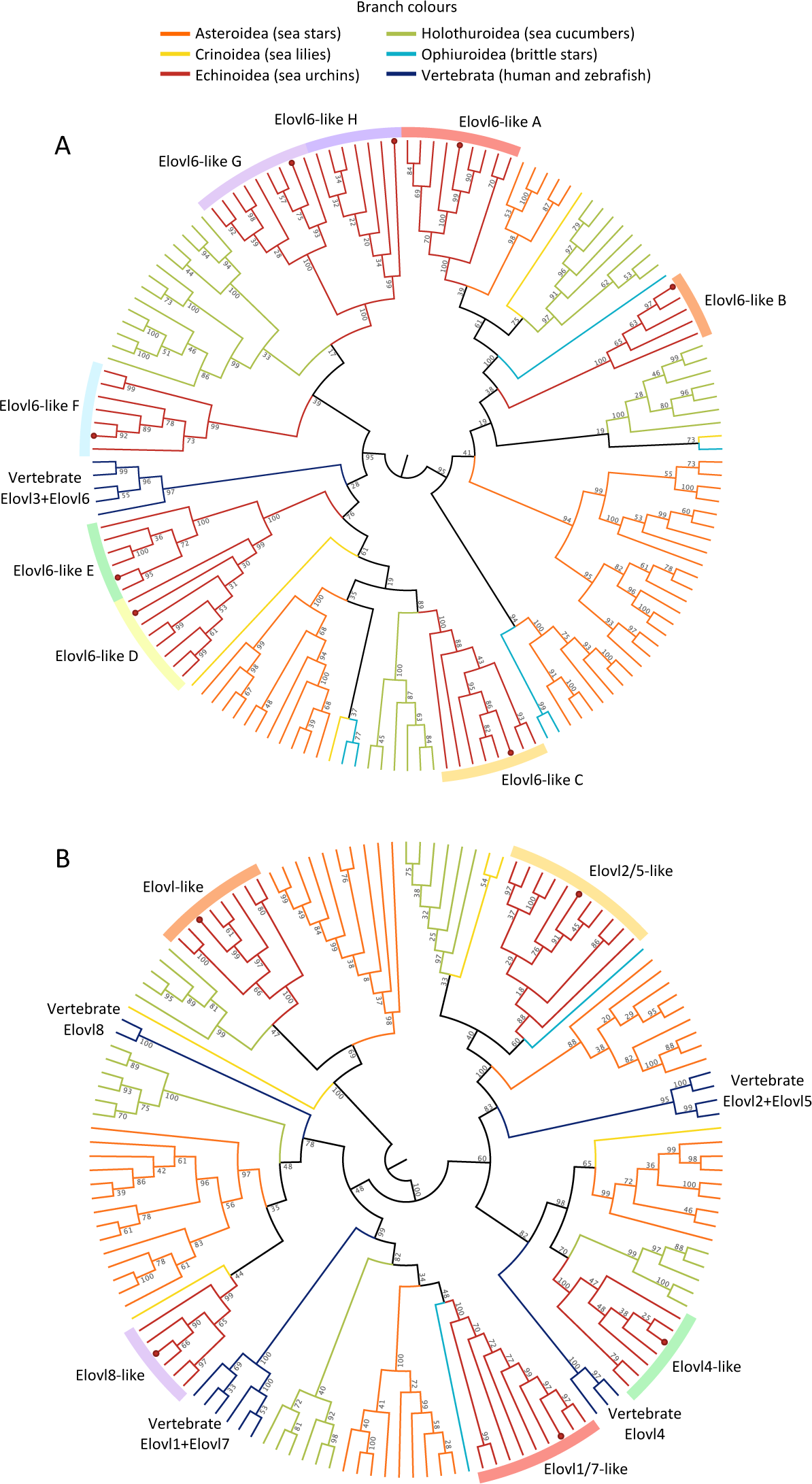
Circular cladograms of the maximum likelihood phylogenetic inference of S/MUFA Elovl (A) and PUFA Elovl (B) from echinoderm species. *H. pulcherrimus* sequences are indicated by red dots in each leaf node. The fully labelled phylogram trees are provided in Supplementary Fig. 2 (S/MUFA Elovl) and Supplementary Fig. 3 (PUFA Elovl).

**Fig. 2.**
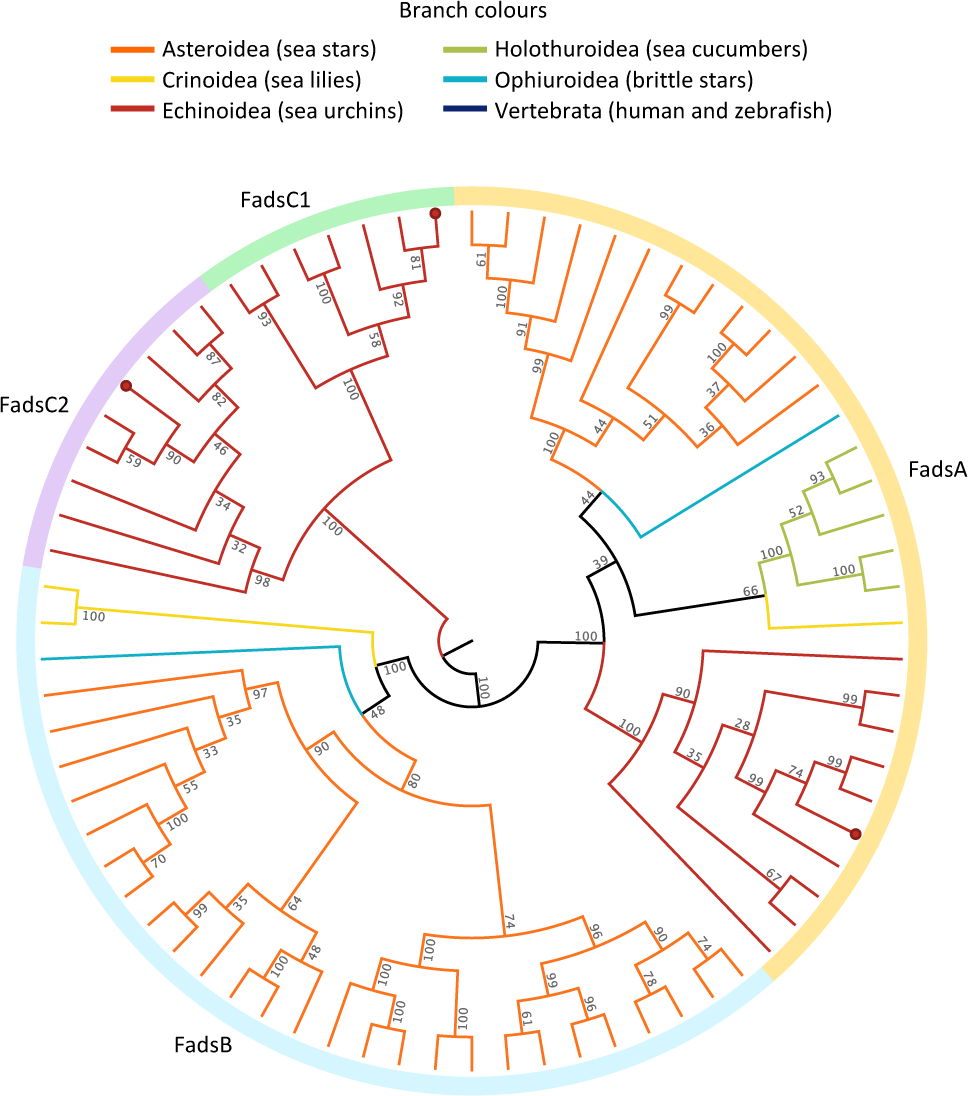
Circular cladogram of the maximum likelihood phylogenetic inference of Fads from echinoderm species. *H. pulcherrimus* sequences are indicated by red dots in each leaf node. The fully labelled tree is provided in Supplementary Fig. 4.

### 3.2. Functional characterisation of H. pulcherrimus Elovl

Regarding S/MUFA Elovl, seven out of eight Elovl showed almost undetectable level of activity towards all substrates tested (Table 1). However, surprisingly, along with an elongase capacity towards the yeast endogenous MUFA (i.e., 16:1n-7 and 18:1n-9) to produce 18:1n-7+20:1n-7 and 20:1n-9, respectively (Fig. 3A, Supplementary Table 2), prominent conversions towards all C_18_ PUFA substrates were observed in the yeast expressing *elovl6-like C* (Fig. 3B and 3C). Table 2 showed substrate conversions (%) of all PUFA Elovl isolated from *H. pulcherrimus*. All PUFA Elovl possessed the elongation capacity towards a series of C_18_ and C_20_ PUFA substrates to produce corresponding +C_2_ products. However, most of conversions are overall low (below 5%) except Elovl1/7-like showing clear elongations towards ARA (20:4n-6) and EPA (20:5n-3) (Table 2). The Elovl1/7-like further showed the activity towards C_22_ PUFA substrates to produce the corresponding C_24_ products (i.e., 24:4n-6 and 24:5n-3) (Table 2).

**Fig. 3.**
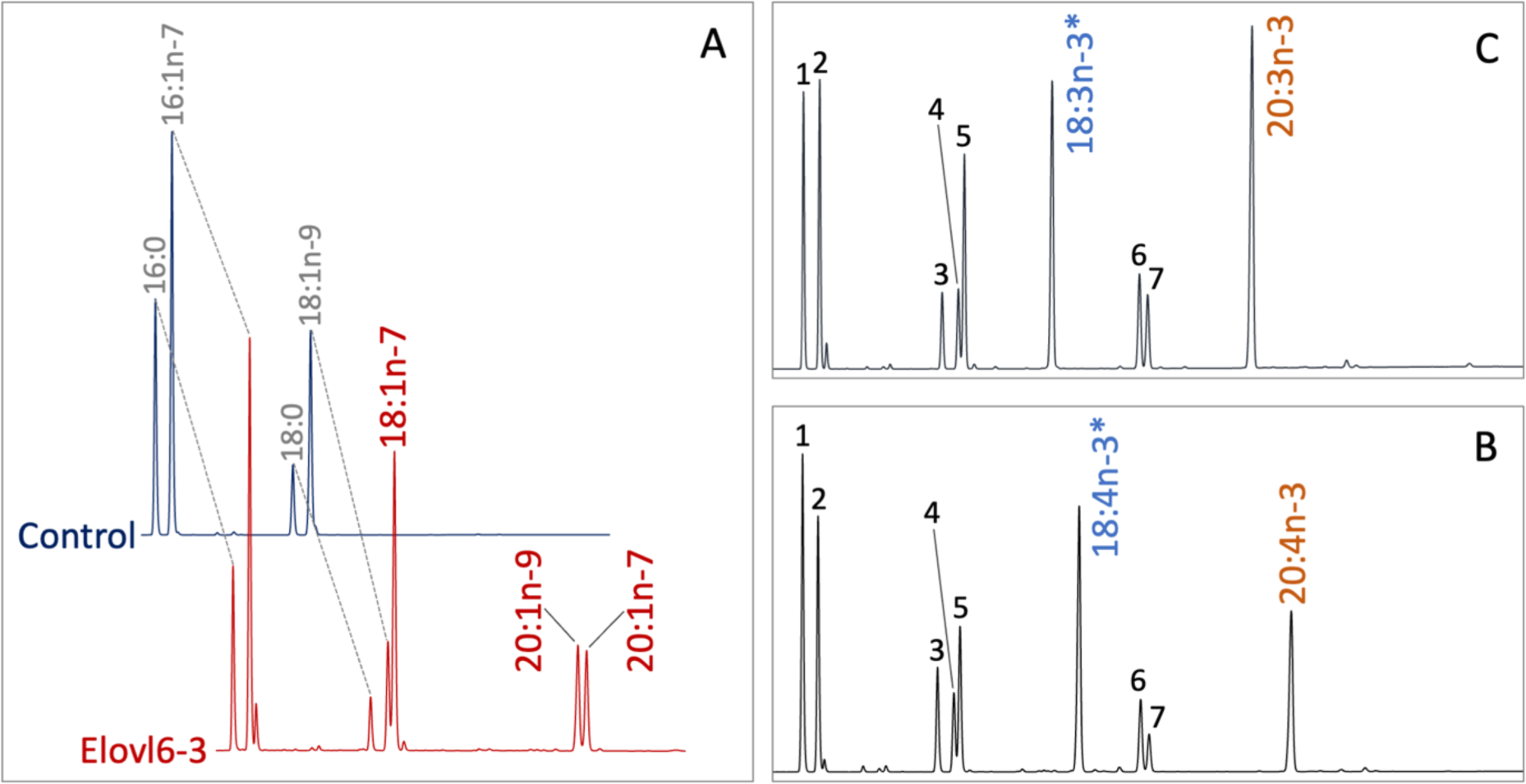
Elovl6-like C isolated from *H. pulcherrimus* showed elongase activities towards various C_18_ MUFA and PUFA to produce the corresponding C_20_ products. (A) GC chromatograms of FAME prepared from the transgenic yeast transformed with an empty pYES2 (Control) or pYES2 containing the full-length ORF of *H. pulcherrimus elovl6-like c*. (B and C) GC chromatograms of FAME prepared from the transgenic yeast transformed with pYES2 containing the full-length ORF of *H. pulcherrimus elovl6-like c* and grown in the presence of 18:2n-6 (B) and 18:3n-3 (C) indicated with an asterisk (*). Fatty acids are numbered as follows: 16:0 (1), 16:1n-7 (2), 18:0 (3), 18:1n-9 (4), 18:1n-7 (5), 20:1n-9 (6) and 20:1n-7 (7).

**Table 1.**
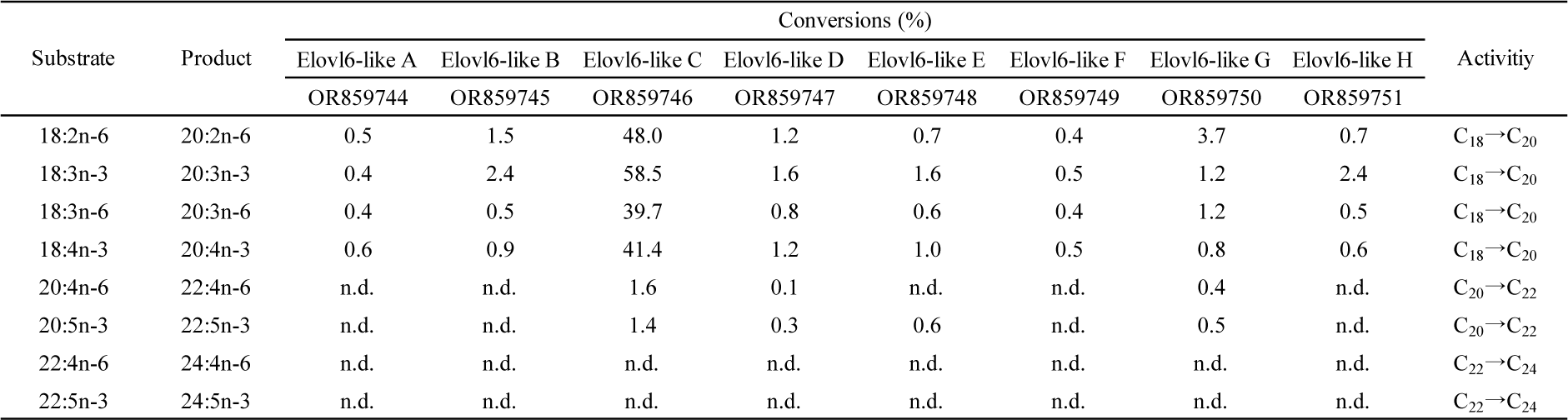
Substrate conversion of the yeast *S. cerevisiae* transformed with pYES2 containing the ORF of the *H. pulcherrimus* elongases belong to S/MUFA subfamily defined by Hashimoto et al. (2008).

**Table 2.** Substrate conversion of the yeast *S. cerevisiae* transformed with pYES2 containing the ORF of the *H. pulcherrimus* elongases belong to PUFA subfamily defined by Hashimoto et al. (2008).

| Substrate | Product | Conversions (%) |  |  |  |  | Activity |
| --- | --- | --- | --- | --- | --- | --- | --- |
|  |  | Elovl-like | Elovl-like | Elovl1/7-like | Elovl4-like | Elovl8-like |  |
|  |  | OR753998 | OR753999 | OR754000 | OR754001 | OR754002 |  |
| 18:2n-6 | 20:2n-6 | 1.6 | 1.1 | 3.4 | 0.6 | 1.5 | C<sub>18</sub>→C<sub>20</sub> |
| 18:3n-3 | 20:3n-3 | 2.5 | 0.2 | 1.9 | 0.7 | 3.9 | C<sub>18</sub>→C<sub>20</sub> |
| 18:3n-6 | 20:3n-6 | 3.4 | 0.6 | 8.6 | 0.7 | 0.8 | C<sub>18</sub>→C<sub>20</sub> |
| 18:4n-3 | 20:4n-3 | 5.7 | 0.4 | 3.0 | 0.8 | 0.7 | C<sub>18</sub>→C<sub>20</sub> |
| 20:4n-6 | 22:4n-6 | 3.1 | 0.2 | 21.7 | 0.4 | 2.0 | C<sub>20</sub>→C<sub>22</sub> |
| 20:5n-3 | 22:5n-3 | 3.7 | 0.4 | 11.6 | 0.9 | 1.7 | C<sub>20</sub>→C<sub>22</sub> |
| 22:4n-6 | 24:4n-6 | n.d. | n.d. | 9.5 | n.d. | n.d. | C<sub>22</sub>→C<sub>24</sub> |
| 22:5n-3 | 24:5n-3 | n.d. | n.d. | 14.7 | n.d. | n.d. | C<sub>22</sub>→C<sub>24</sub> |

### 3.3. Functional characterization of H. pulcherrimus Fads

As shown in Table 3, clear product peaks (ARA and EPA) observed in the transgenic yeast expressing *H. pulcherrimus* FadsA grown in the presence of 20:3n-6 and 20:4n-3, respectively, denoting that FadsA is a Δ5 desaturase (Table 3). The Δ5 desaturation by FadsA can also be happened onto 20:2n-6 and 20:3n-3 to produce NMI-FA 20:3^Δ5,11,14^ and 20:4^Δ5,11,14,17^, respectively. In addition, the transgenic yeast expressing FadsA was able to desaturate MUFA substrates, namely 20:1n-9 and 20:1n-7 into the NMI-FA 20:2^Δ5,11^ and 20:2^Δ5,13^, respectively. The mass spectra of DMOX derivative of all those NMI-FA contained a diagnostic ion for Δ5 desaturation at m/z = 153 (Supplementary Fig. 5 and 6). The 20:2^Δ5,11^ and 20:2^Δ5,13^ were clearly distinguished by the position of the characteristic gaps of 12 atomic mass units (amu) between m/z = 222 and 234, and m/z = 250 and 262, respectively in their mass spectra (Supplementary Fig. 5). The MS profiles of 20:3^Δ5,11,14^ and 20:4^Δ5,11,14,17^ also exhibited the characteristic 12 amu gaps indicating the double bonds at Δ11,14 and Δ11,14,17 respectively (Supplementary Fig. 6). Unlike FadsA, FadsC2 showed clear Δ8 desaturase activity towards 20:2n-6 and 20:3n-3 to produce 20:3n-6 and 20:4n-3, respectively (Table 3), while only trace level of activities towards 20:3n-3 and 20:3n-6 were detected in the yeast expressing FadsC1 (Table 3).

**Table 3.** Substrate conversion of the yeast *S. cerevisiae* transformed with pYES2 containing the ORF of the *H. pulcherrimus* fatty acyl desaturases.

| Substrate | Product | Conversion (%) |  |  | Activity |
| --- | --- | --- | --- | --- | --- |
|  |  | FadsA | FadsC1 | FadsC2 |  |
|  |  | OR545538 | OR545539 | OR545540 |  |
| 18:2n-6 | 18:3n-6 | n.d. | n.d. | 0.5 | Δ6 |
| 18:3n-3 | 18:4n-3 | n.d. | n.d. | 0.1 | Δ6 |
| 20:2n-6 | 20:3n-6 | n.d. | n.d. | 40.5 | Δ8 |
| 20:3n-3 | 20:4n-3 | n.d. | 0.2 | 11.7 | Δ8 |
| 20:3n-6 | 20:4n-6 | 50.2 | n.d. | n.d. | Δ5 |
| 20:4n-3 | 20:5n-3 | 34.7 | n.d. | n.d. | Δ5 |
| 22:4n-6 | 22:5n-6 | n.d. | n.d. | n.d. | Δ4 |
| 22:5n-3 | 22:6n-3 | n.d. | n.d. | n.d. | Δ4 |

**Table 4.**
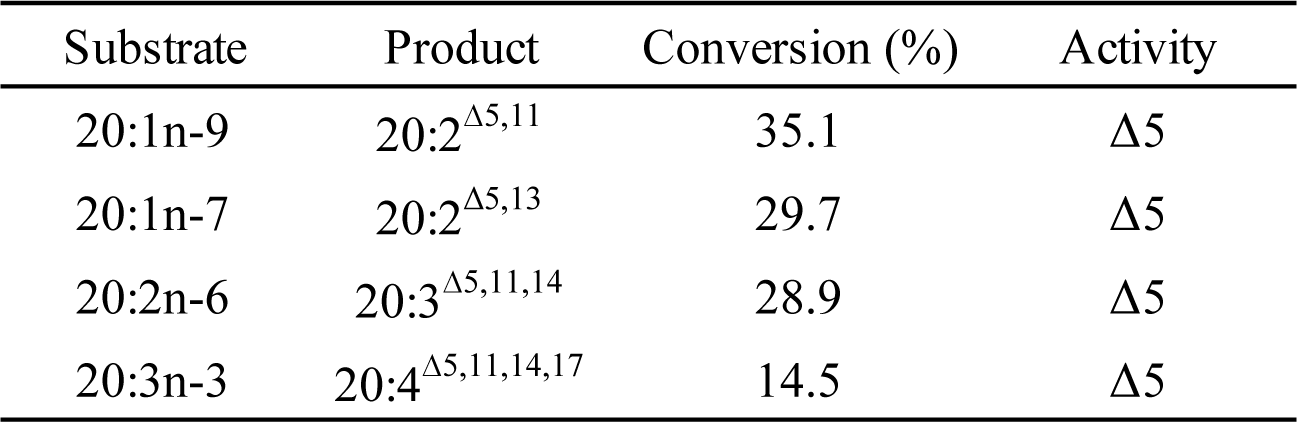
Δ5 desaturase activity towards various C_20_ substrates to produce NMI-FA in the transgenic yeast expressing *H. pulcherrimus* FadsA.

## 4. Discussion

In marine organisms, LC-PUFA has important roles in several physiological processes such as reproduction (Sargent et al., 1999; Furuita et al., 2002), brain and eye development (Furuita et al., 1998; Benítez-Santana et al., 2007) and growth performance (Skalli & Robin, 2004). Studies have shown that dietary LC-PUFA can significantly promote growth in juvenile sea urchins, including *P. lividus* (Liu et al., 2007). However, other research indicates that sea urchins can achieve normal development and growth without dietary LC-PUFA (Fernandez & Boudouresque, 2000; Liyana-Pathirana et al., 2002; Vizzini et al., 2015). For example, although ARA enhances gonad development, nutritional value and growth performance in adult *S. intermedius* (Zuo et al., 2018), DHA appears non-essential as its deficiency does not affect gametogenesis or survival (Carboni et al., 2013; Zuo et al., 2018). The content of DHA in sea urchins remains very low on a DHA-deficient diet, and the absence of ARA in the diet does not affect its levels in sea urchins, when 18:2n-6 was provided in their diet. Similarly, the levels of EPA in sea urchins are unaffected by diets lacking EPA but containing 18:3n-3 (Carboni et al., 2013; González-Durán et al., 2008). The present study unequivocally demonstrates that the sea urchin, *H. pulcherrimus* possesses a complete set of enzymes necessary to synthesise ARA and EPA endogenously from their corresponding C_18_ PUFA precursors, 18:2n-6 and 18:3n-3, respectively. This finding underlines the importance of including these C_18_ precursors in the sea urchin diet to facilitate the synthesis of ARA and EPA. Furthermore, the presence of multiple copies of potential orthologues of vertebrate Elovl6, characterised as a MUFA elongase, were found in *H. pulcherrimus*. One of these, Elovl6-like C), appears to play a prominent role in the LC-PUFA biosynthesis. Functional analysis of these enzymes also allowed us to delineate C_20_ NMI-FA biosynthetic pathways in *H. pulcherrimus*. This analysis demonstrated that, through the action of enzymes isolated in the present study, including FadsA with Δ5 desaturase activity and several elongases like Elovl6-like C, *H. pulcherrimus* is capable of synthesising a range of C_20_ methylene-interrupted PUFA (20:2n-6, 20:3n-3, 20:3n-6, 20:4n-3, ARA and EPA) and four C_20_ NMI-FA (20:2^Δ5,11^, 20:2^Δ5,13^, 20:3^Δ5,11,14^, 20:4^Δ5,11,14,17^) from C_18_ precursors (Fig. 5).

**Fig. 4.**
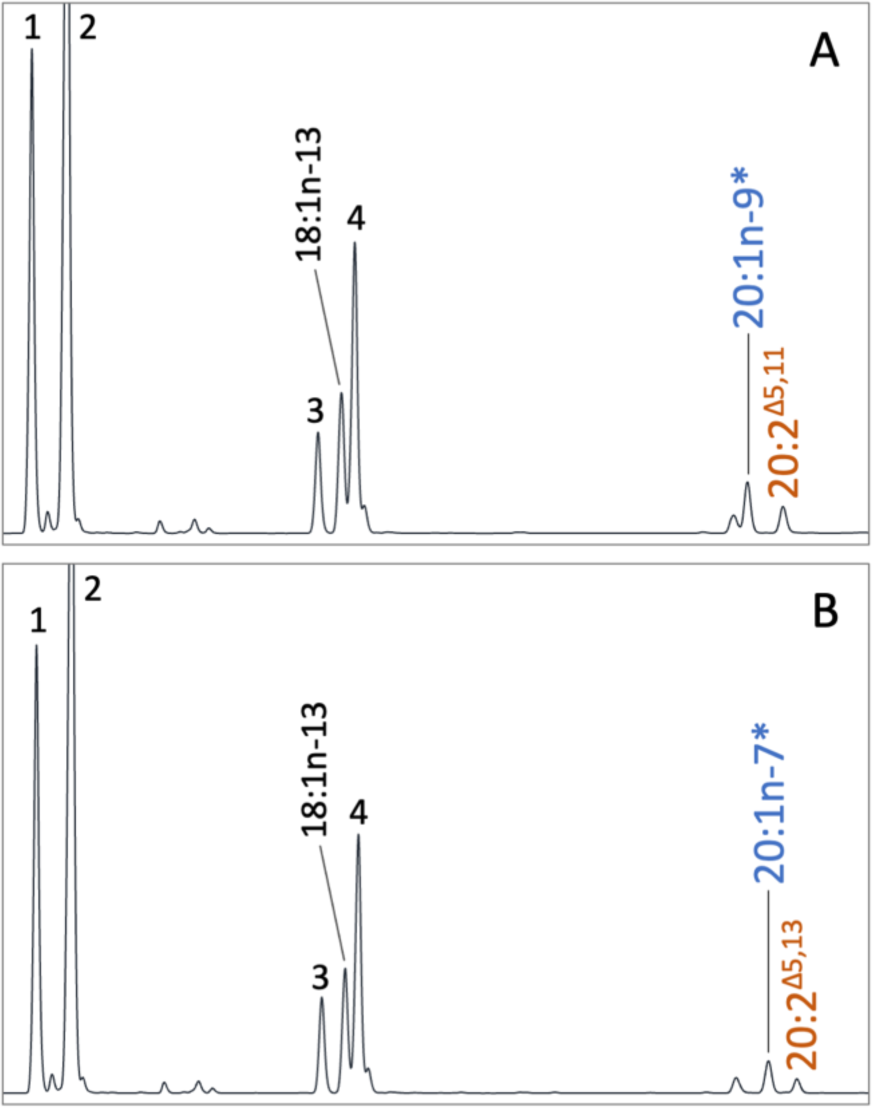
*H. pulcherrimu*s FadsA enables production of C_20_ non-methylene interrupted dienoic fatty acids. GC chromatograms of FAME prepared from the transgenic yeast transformed with pYES2 containing the full-length ORF of *H. pulcherrimus fadsa* and grown in the presence of 20:1n-9 (A) and 20:1n-7 (C) indicated with an asterisk (*). 18:1n-13 is a Δ5 desaturated product from 18:0. The yeast endogenous fatty acids are numbered as follows: 16:0 (1), 16:1n-7 (2), 18:0 (3), 18:1n-9 (4).

**Fig. 5.**
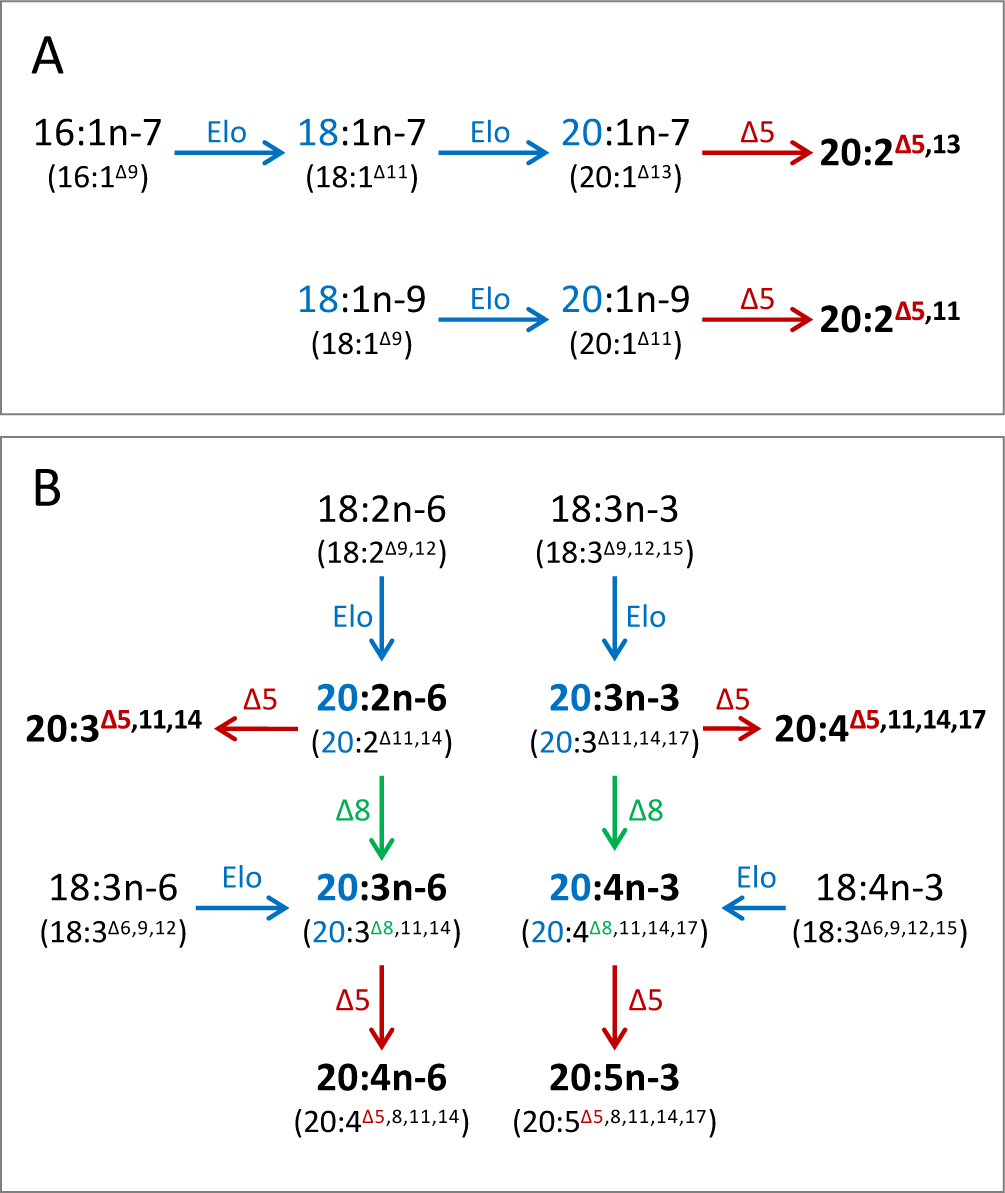
Biosynthetic pathways of various C_20_ PUFA in *H. pulcherrimus*. (A) The biosynthetic pathways of C_20_ dienoic NMI-FA from C_16_ and C_18_ MUFA substrates. (B) The biosynthetic pathways of various C_20_ PUFA including physiologically important arachidonic acid (ARA, 20:4n-6) and eicosapentaenoic acid (EPA, 20:5n-3) as well as trienoic and tetraenoic NMI-FA (20:3^Δ5,11,14^ and 20:4^Δ5,11,14,17^, respectively).

The comprehensive retrieval of Elovl enzymes from sea urchins revealed a marked increase in the copy number of *elovl6-like* gene not only in *H. pulcherrimu*s but also in other sea urchin species. In the ML phylogenetic analysis, each sea urchin Elovl6-like (A to H) formed a well-supported clade (Fig. 1A), suggesting similar PUFA elongase activity across these variants as observed in *H. pulcherrimu*s. However, the phylogenetic tree resolution was not sufficiently high to fully determine the presence, absence, and relationships of each Elovl6-like orthologue among different echinoderms. Interestingly, a clade containing several sea cucumber sequences with unknown function appeared to be a direct sister group to the sea urchin Elovl6-like C. Given the close relation between sea urchins sea cucumbers (Telford et al., 2014; Reich et al., 2015), it is likely that *elovl6-like C* gene emerged in their common ancestor. Although mammalian Elovl6 has been characterised as an elongase showing activity towards SFA and MUFA, none of the Elovl6 in *H. pulcherrimus* exhibited an elongase activity towards the yeast endogenous MUFA (16:1n-7 and 18:1n-9) except for Elovl6-like C. This enzyme was capable of elongating several C_18_ unsaturated fatty acid substrates regardless of their degree of unsaturation, making it potentially the most physiologically significant among the Elovl6-like enzymes isolated.

Regarding PUFA elongases, four Elovl from *H. pulcherrimus* (Elovl2/5-like, Elovl4-like, Elovl1/7-like and Elovl8-like) clustered with corresponding vertebrate Elovl enzymes in physogenetic analysis (Fig. 1B). However, there were no exact one-to-one orthologous enzymes for Elovl2 and Elovl5, as well as Elovl1 and Elovl7, likely owing to gene duplications in ancestral vertebrates or later-diverging deuterostomes (Monroig et al., 2016). Unlike these four, one Elovl from *H. pulcherrimus* did not cluster with any vertebrate Elovl and is likely widespread among echinoderm species, including sea stars, sea cucumbers and one sea lily. Most Elovl showed elongase activity towards multiple C_18_ and C_20_ PUFA substrates, but only Elovl1/7-like was capable of elongating C_22_ PUFA substrates to produce C_24_ products (24:4n-6 and 24:5n-3), which, however, to the best of our knowledge, have not been detected in the sea urchin tissues, suggesting limited biological significance. Indeed, another C_24_ PUFA, 24:6n-3 is uncommon in sea urchin species, albeit the presence of a certain level of C_24_ PUFA in some other echinoderms (brittle stars and sea lilies) (Zhukova, 2022).

The regioselectivity detected in three Fads from *H. pulcherrimus* are fairly consistent with those of previous studies in *P. lividus* (Kabeya et al., 2017); one FadsA with a Δ5 desaturase activity and two FadsC with either overall low activity (FadsC1) or a Δ8 desaturase activity (FadsC2). In addition to the previously reported Δ5 desaturase activity towards 20:2n-6 and 20:3n-3 to produce NMI-FA, 20:3^Δ5,11,14^ and 20:4^Δ5,11,14,17^, respectively (Kabeya et al. 2017), FadsA from *H. pulcherrimus* was able to introduce Δ5 desaturation in two MUFA substrates 20:1n-9 and 20:1n-7 to produce 20:2^Δ5,11^ and 20:2^Δ5,13^, respectively. Since sea urchins generally contain significant levels of those dienoic NMI-FA (Zhukova, 2022), this activity would be a common feature of FadsA in sea urchins. Furthermore, Given the FadsA was widespread among all echinoderm classes and Δ5 desaturated NMI-FA are commonly found in not only sea urchins but also other echinoderms such as sea cucumbers and sea stars (Kabeya et al., 2017; Zhukova, 2022), it could be speculated that those Δ5 desaturated NMI-FA were endogenously synthesised and accumulated in their body, despite unknown physiological functions. Unlike FadsA, distributions of FadsB and FadsC were more class-specific; FadsB is present in sea star, brittle star and sea lily species, and FadsC (FadsC1 and FadsC2) are present exclusively in sea urchin species. Considering the phylogenetic relationships of echinoderms (Telford et al. 2014; Reich et al. 2015), *fadsb* gene might be lost in the common ancestor of Echinozoa (sea urchins and sea cucumbers).

In conclusion, the present study established that *H. pulcherrimus* possessed thirteen Elovl and three Fads enzymes potentially involved in PUFA biosynthesis. Notably, Elovl6-like C exhibits elongase activity towards C_18_ PUFA substrates to produce several C_20_ PUFA products, an unexpected function given that Elovl6 is typically categorised as a S/MUFA elongase in vertebrate species. This finding underscores the risk of drawing conclusions about enzymatic functions based solely on amino acid sequences and/or phylogenetic analysis, which could lead to inaccurate interpretations of a species’ biosynthetic capabilities of PUFA. Regarding desaturase functions, Δ5 desaturase (FadsA) and Δ8 desaturase activity (FadsC2) were found in *H. pulcherrimus*. Together with Elovl activities, *H. pulcherrimus* is capable of synthesising physiologically important ARA and EPA from 18:2n-6 and 18:3n-3, respectively *via* the so-called “Δ8 pathway” (Kabeya et al., 2017). In addition, FadsA was able to synthesise several NMI-FA from both C_20_ PUFA and MUFA substrates. The results obtained in the present study provide solid molecular evidence for the dietary essential fatty acids in the sea urchin and endogenous production of particularly unusual fatty acids, NMI-FA.

## Acknowledgments

YP is supported by the International Education for Marine Environment and Energy Use - Japan-China-Korea Exchange Program- (JCK Program) at Tokyo University of Marine Science and Technology.

## Author’s contributions

NK initially conceived the study and design experiments. YP carried out isolation and functional characterisation of genes. Phylogenetic analyses were performed by YP and NK. NK and YH interpreted the results obtained from fatty acid analysis with YP. YP prepared the first draft of the manuscript and edited with NK and YH. The manuscript was finalised by all authors.

## Availability of data and materials

The full-length ORF sequences isolated from *H. pulcherrimus* in the present study were deposited into the NCBI GenBank; S/MUFA elongases (accessions OR859744-OR859751), PUFA elongases (OR753998-OR754002) and fatty acid desaturases (OR545538-OR545540).

## References

Archana A, Babu KR. 2016 Nutrient composition and antioxidant activity of gonads of sea urchin *Stomopneustes variolaris*. Food Chem. 197(Pt A), 597–602. (doi: 10.1016/j.foodchem.2015.11.003)

Benítez-Santana T, Masuda R, Carrillo EJ, Ganuza E, Valencia A, Hernández-Cruz CM, Izquierdo MS. 2007 Dietary n-3 HUFA deficiency induces a reduced visual response in gilthead seabream *Sparus aurata* larvae. Aquaculture. 264(1-4): 408–417. (doi: 10.1016/j.aquaculture.2006.10.024)

Brouwer IA, Geenlen A, Katan MB. 2006 n-3 Fatty acids, cardiac arrhythmia and fatal coronary heart disease. Prog Lipid Res. 45, 357–367. (doi:10.1016/j.plipres.2006.02.004)

Castilla-Gavilán M, Buzin F, Cognie B, Dumay J, Turpin V, Decottignies P. 2018 Optimising microalgae diets in sea urchin *Paracentrotus lividus* larviculture to promote aquaculture diversification. Aquaculture. 490, 251–259. (doi:10.1016/j.aquaculture.2018.02.003)

Carboni S, Vignier J, Chiantore M, Tocher DR, Migaud, H. 2012 Effects of dietary microalgae on growth, survival and fatty acid composition of sea urchin *Paracentrotus lividus* throughout larval development. Aquaculture. 324-325, 250–258. (doi:10.1016/j.aquaculture.2011.10.037)

Carboni S, Hughes AD, Atack T, Tocher DR, Migaud H. 2013 Fatty acid profiles during gametogenesis in sea urchin (*Paracentrotus lividus*): effects of dietary inputs on gonad, egg and embryo profiles. Comp Biochem Physiol A Mol Integr Physiol. 164, 376–82. (doi: 10.1016/j.cbpa.2012.11.010)

Capella-Gutierrez S, Silla-Martinez JM, Gabaldón T. 2009 trimAl: a tool for automated alignment trimming in large-scale phylogenetic analyses. Bioinformatics 25, 1972 1973. (doi:10.1093/bioinformatics/btp348)

Darriba D, Posada D, Kozlov AM, Stamatakis A, Morel B, Flouri T. 2019 ModelTest-NG: a new and scalable tool for the selection of DNA and protein evolutionary models. Mol Biol Evol 37, 291–294. (doi:10.1093/molbev/msz189)

Fernandez C, Boudouresque CF. 2000 Nutrition of the sea urchin *Paracentrotus Lividus* (Echinodermata: Echinoidea) fed different artificial food. Mar Ecol Prog Ser. 204, 131–41. (doi:10.3354/meps204131)

Furuita H, Takeuchi T, Uematsu K. 1998 Effects of eicosapentaenoic and docosahexaenoic acids on growth, survival and brain development of larval Japanese flounder (*Paralichthys olivaceus*). Aquaculture. 161, 269–279. (doi: 10.1016/S0044-8486(97)00275-5)

Furuita H, Tanaka H, Yamamoto T, Suzuki N, Takeuchi T. 2002 Effects of high levels of n− 3 HUFA in broodstock diet on egg quality and egg fatty acid composition of Japanese flounder, *Paralichthys olivaceus*. Aquaculture. 210, 323–333. (doi: 10.1016/S0044-8486(01)00855-9)

González-Durán E, Castell JD, Robinson SMC, Blair TJ. 2008 Effects of dietary lipids on the fatty acid composition and lipid metabolism of the green sea urchin *Strongylocentrotus droebachiensis*. Aquaculture. 276, 120–129. (doi: 10.1016/j.aquaculture.2008.01.010)

Hashimoto K, Yoshizawa AC, Okuda S, Kuma K, Goto S, Kanehisa M. 2008 The repertoire of desaturases and elongases reveals fatty acid variations in 56 eukaryotic genomes. J Lipid Res. 49,183–191. (doi:10.1194/jlr.M700377-JLR200)

Kabeya N, Fonseca MM, Ferrier DEK, Navarro JC, Bay LK, Francis DS, Tocher DR, Castro LFC, Monroig Ó. 2018 Genes for de novo biosynthesis of omega-3 polyunsaturated fatty acids are widespread in animals. Sci Adv. 4, eaar6849. (doi:10.1126/sciadv.aar6849)

Kabeya N, Ogino M, Ushio H, Haga Y, Satoh S, Navarro JC, Monroig Ó. 2021 A complete enzymatic capacity for biosynthesis of docosahexaenoic acid (DHA, 22: 6n–3) exists in the marine Harpacticoida copepod Tigriopus californicus. Open Biol. 11, 200402. (doi: 10.1098/rsob.200402)

Kabeya N, Sanz-Jorquera A, Carboni S, Davie A, Oboh A, Monroig O. 2017 Biosynthesis of polyunsaturated fatty acids in sea urchins: Molecular and functional characterisation of three fatty acyl desaturases from *Paracentrotus lividus* (Lamark 1816). PlOS ONE. 12, e0169374. (doi: 10.1371/journal.pone.0169374)

Katoh K, Rozewicki J, Yamada KD. 2017 MAFFT online service: multiple sequence alignment, interactive sequence choice and visualization. Brief Bioinform 30, bbx108-. (doi:10.1093/bib/bbx108)

Kelly JR, Scheibling RE, Iverson SJ, Gagnon P. 2008 Fatty acid profiles in the gonads of the sea urchin *Strongylocentrotus droebachiensis* on natural algal diets. Mar. Ecol. Prog. Ser. 373, 1–9. (doi: 10.3354/meps07746)

Kozlov AM, Darriba D, Flouri T, Morel B, Stamatakis A. 2019 RAxML-NG: A fast, scalable, and user-friendly tool for maximum likelihood phylogenetic inference. Bioinformatics 35, 4453–4455. (doi:10.1093/bioinformatics/btz305)

Liyana-Pathirana C, Shahidi F, Whittick A, Hooper R. 2002 Lipid and lipid soluble components of gonads of green sea urchin (*Strongylocentrotus droebachiensis*). J Food Lipids. 9, 105–126. (doi: 10.1111/j.1745-4522.2002.tb00213.x)

Liu H, Kelly MS, Cook EJ, Black K, Orr H, Zhu JX, Dong SL. 2007 The effect of diet type on growth and fatty-acid composition of sea urchin larvae, I. *Paracentrotus lividus* (Lamarck, 1816) (Echinodermata). Aquaculture. 264. 247–262. (doi:10.1016/j.aquaculture.2006.12.021)

Miyashita K, Hosokawa M. 2010 Future direction of lipid nutrition and development of functional foods - Gift from seaweeds and its application. Journal of Lipid Nutrition. 19, 39–45. (doi:10.4010/jln.19.39)

Monroig Ó, Kabeya N. 2018 Desaturases and elongases involved in polyunsaturated fatty acid biosynthesis in aquatic invertebrates: a comprehensive review. Fish Sci. 84, 911–928. (doi: 10.1007/s12562-018-1254-x)

Monroig Ó, Lopes-Marques M, Navarro JC, Hontoria F, Ruivo R, Santos MM, Venkatesh B, Tocher DR, Castro LF. 2016 Evolutionary functional elaboration of the Elovl2/5 gene family in chordates. Sci Rep. 6, 20510. (doi: 10.1038/srep20510).

Paradis M, Ackman RG. 1977 Potential for employing the distribution of anomalous non-methylene-interrupted dienoic fatty acids in several marine invertebrates as part of food web studies. Lipids. 12, 170–176. (doi: 10.1007/BF02533289)

Rahman MA, Arshad A, Yusoff FM. 2014 Sea urchins (Echinodermata:Echinoidea): Their biology, culture and bioactive compounds. International Conference on Aqricultural, Ecological and Medical Sciences (AEMS-2014) July 3-4, 2014 London (United Kingdom). (doi:10.15242/iicbe.c714075)

Reich A, Dunn C, Akasaka K, Wessel G. 2015 Phylogenomic analyses of Echinodermata support the sister groups of Asterozoa and Echinozoa. PLoS ONE 10, e0119627. (doi:10.1371/journal.pone.0119627)

Sargent J, Bell G, McEvoy L, Tocher D, Estevez A. 1999 Recent developments in the essential fatty acid nutrition of fish. Aquaculture. 177, 191–199. (doi: 10.1016/S0044-8486(99)00083-6)

Sato D, Ando Y, Tsujimoto R, Kawasaki K. 2001 Identification of novel non-methylene-interrupted fatty acids, 7E,13E-20:2, 7E,13E,17Z-20:3, 9E,15E,19Z-22:3, and 4Z,9E,15E,19Z-22:4, in Ophiuroidea (brittle star) lipids. Lipids. 36, 1371–1375. (doi: 10.1007/s11745-001-0854-x)

Skalli A and Jean HR. 2004 Requirement of n-3 long chain polyunsaturated fatty acids for European sea bass (*Dicentrarchus labrax*) juveniles: growth and fatty acid composition. Aquaculture. 240, 399–415. (doi: 10.1016/j.aquaculture.2004.06.036)

Sun J, Chiang FS. 2015 Use and Exploitation of Sea Urchins. Echnioderm Aquaculture Chapter 2. 25–44. (Online ISBN: 9781119005810)

Takagi T, Kaneniwa M, Itabashi Y, Ackman RG. 1986 Fatty acids in echinoidea: Unusual cis-5-olefinic acids as distinctive lipid components in sea urchins. Lipids. 21, 558–565. (doi: 10.1007/BF02534052)

Telford MJ, Lowe CJ, Cameron CB, Ortega-Martinez O, Aronowicz J, Oliveri P, Copley RR. 2014 Phylogenomic analysis of echinoderm class relationships supports Asterozoa. Proc. R. Soc. B: Biol. Sci. 281, 20140479. (doi:10.1098/rspb.2014.0479)

Vizzini S, Miccichè L, Vaccaro A, Mazzola A. 2015 Use of fresh vegetable discards as sea urchin diet: effect on gonad index and quality. Aquacult Int. 23, 127–139. (doi:10.1007/s10499-014-9803-5)

Yehuda S, Rabinovitz S, Carasso RL, Mostofsky DI. 2002 The role of polyunsaturated fatty acids in restoring the aging neuronal membrane. Neurobiol Aging. 23, 843–53. (doi: 10.1016/s0197-4580(02)00074-x)

Zhukova NV. 2022 Fatty acids of echinoderms: Diversity, current applications and future opportunities. Mar Drugs. 21(1):21. (doi: 10.3390/md21010021).

Zuo R, Li M, Ding J, Chang Y. 2018 Higher dietary arachidonic acid levels improved the growth performance, gonad development, nutritional value, and antioxidant enzyme activities of adult sea urchin (*Strongylocentrotus intermedius*). J. Ocean Univ. China. 17, 932–940. (doi: 10.1007/s11802-018-3606-7)

