## Supplementary_Figures for "Enzymes enabling the biosynthesis of various C_20_ polyunsaturated fatty acids in a sea urchin *Hemicentrotus pulcherrimus*"

### Supplementary Fig. 1: General LC-PUFA biosynthetic pathway

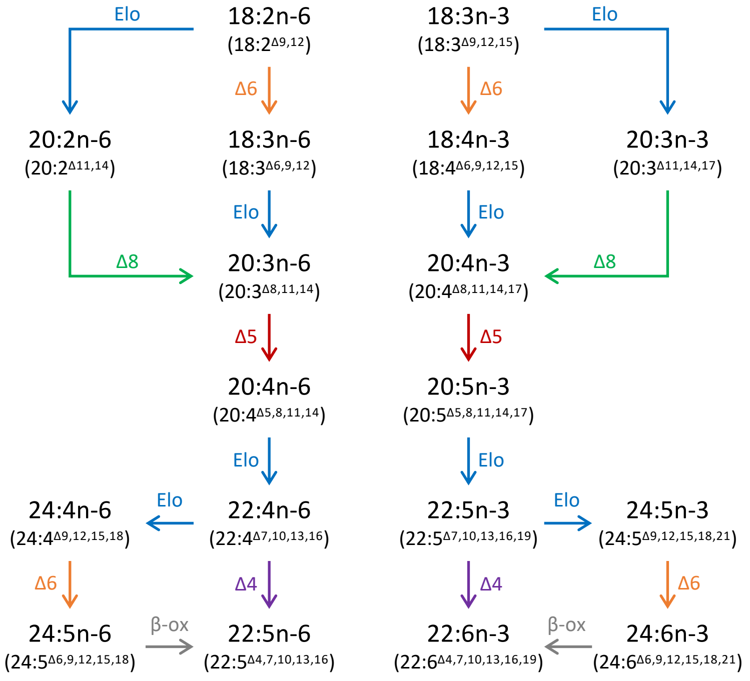

### Supplementary Fig. 2: S/MUFA elongase tree

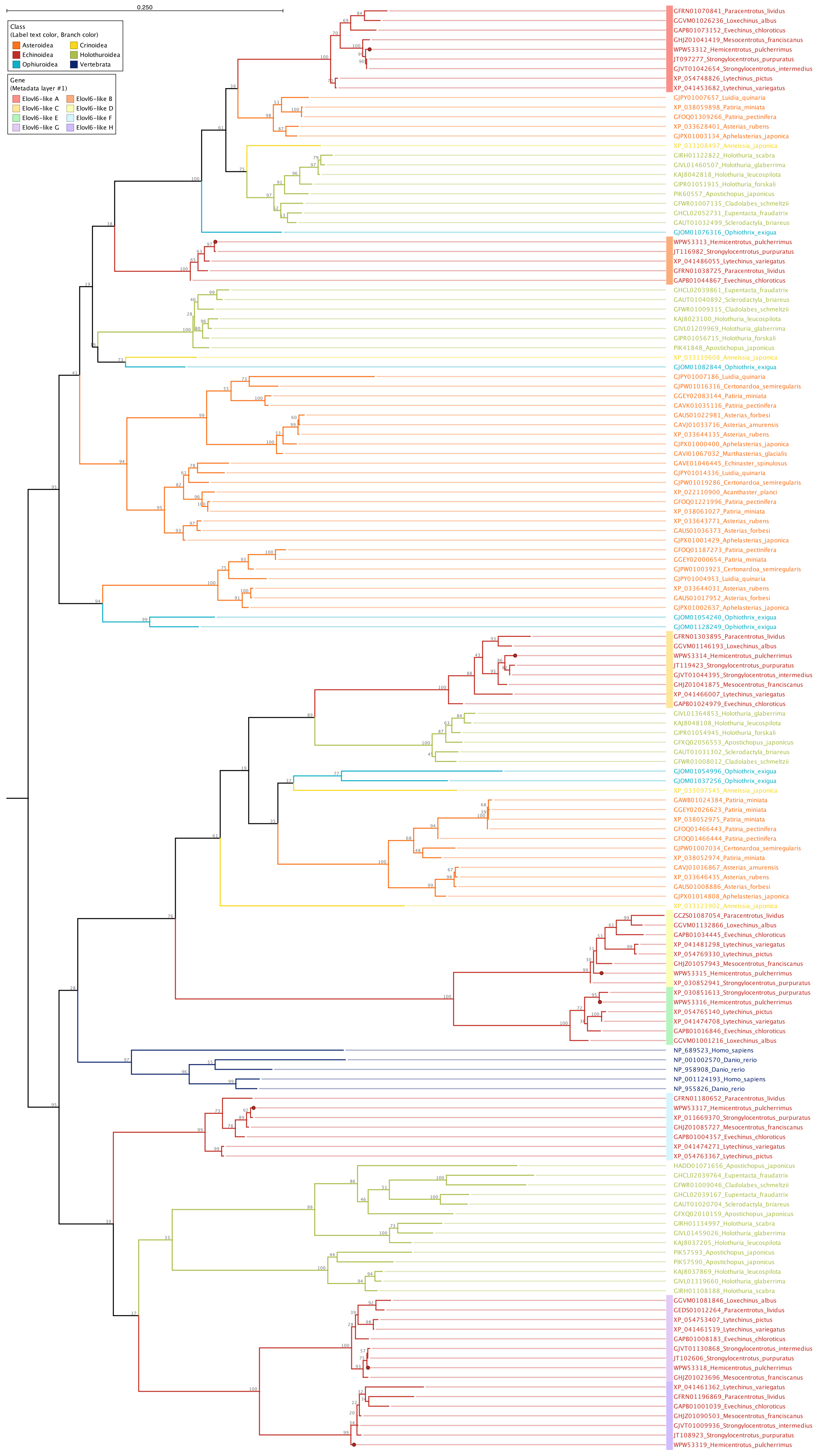

### Supplementary Fig. 3: PUFA elongase tree

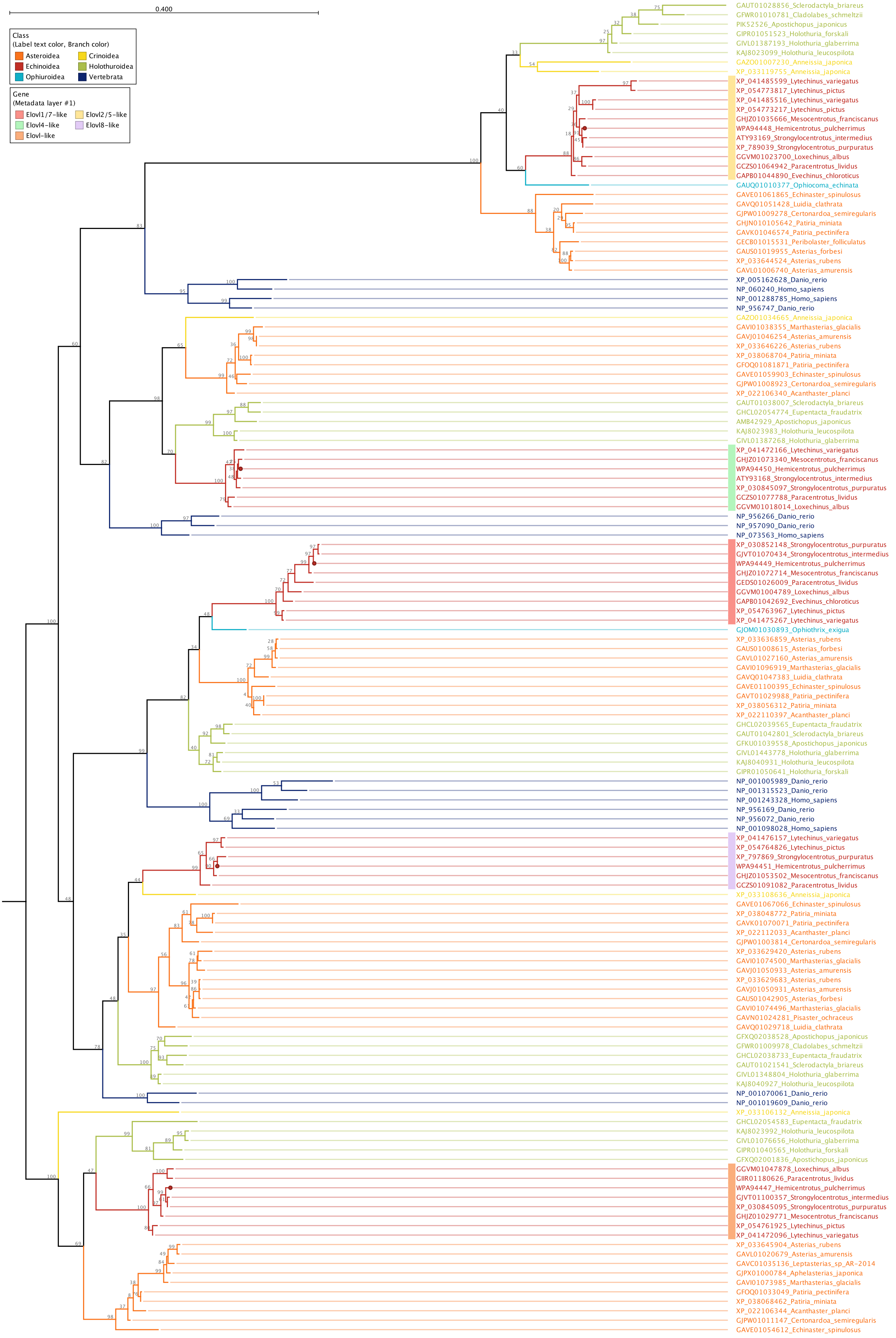

### Supplementary Fig. 4: Desaturase tree

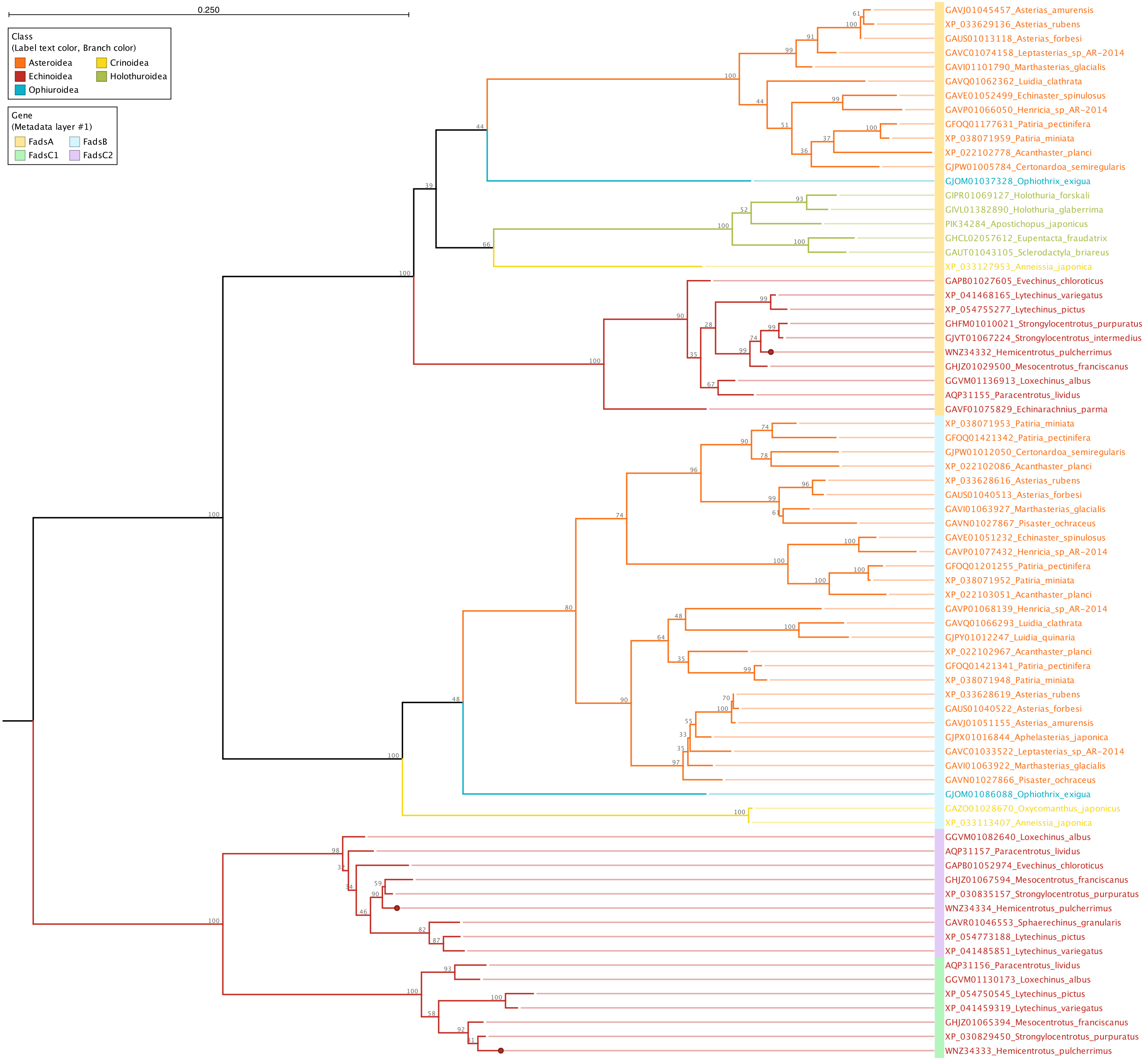

### Supplementary Fig. 5: Mass spectra – NMI dienoic fatty acids

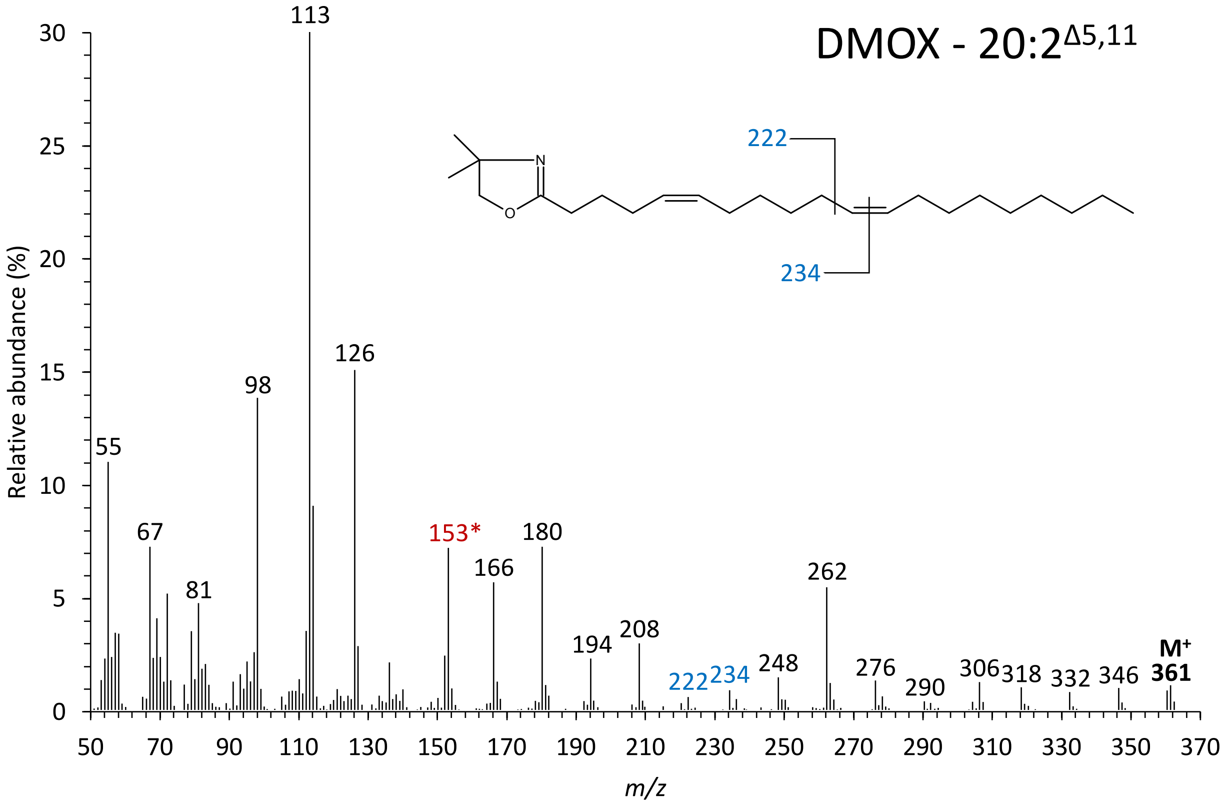


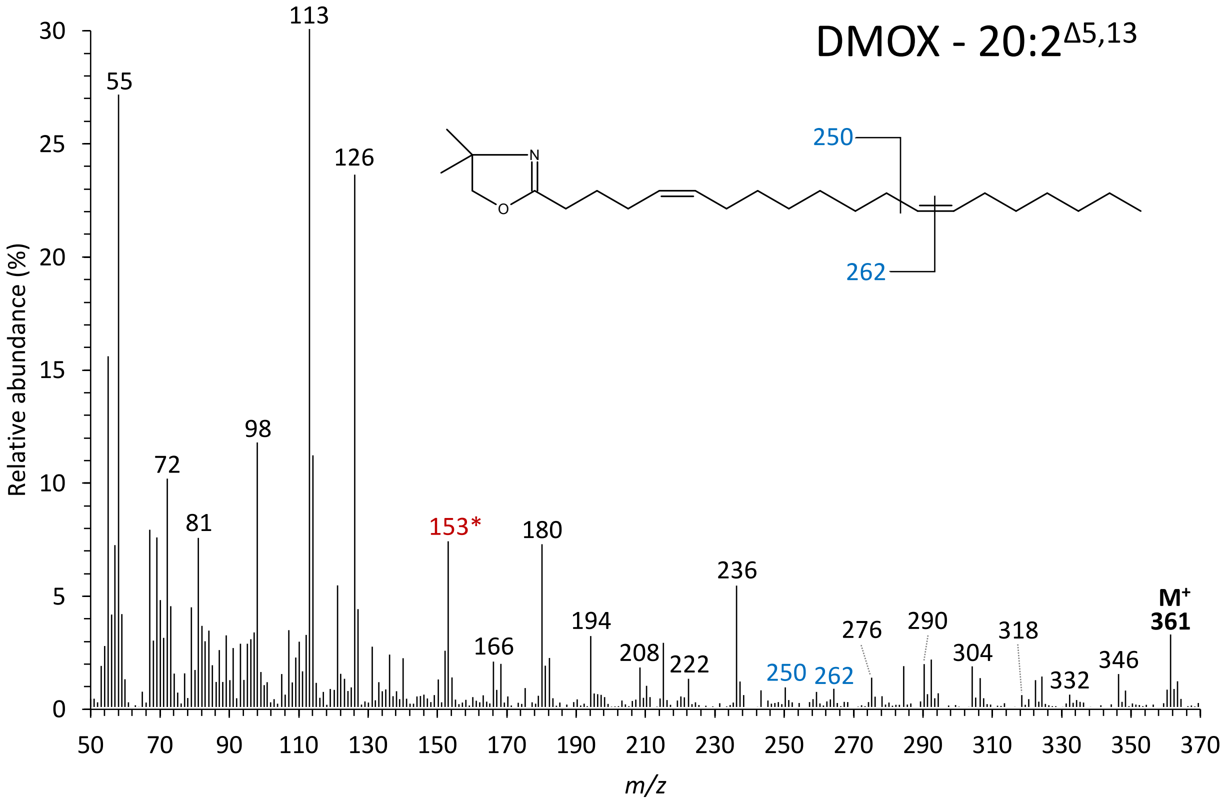

### Supplementary Fig. 6: Mass spectra – NMI polyenoic fatty acids

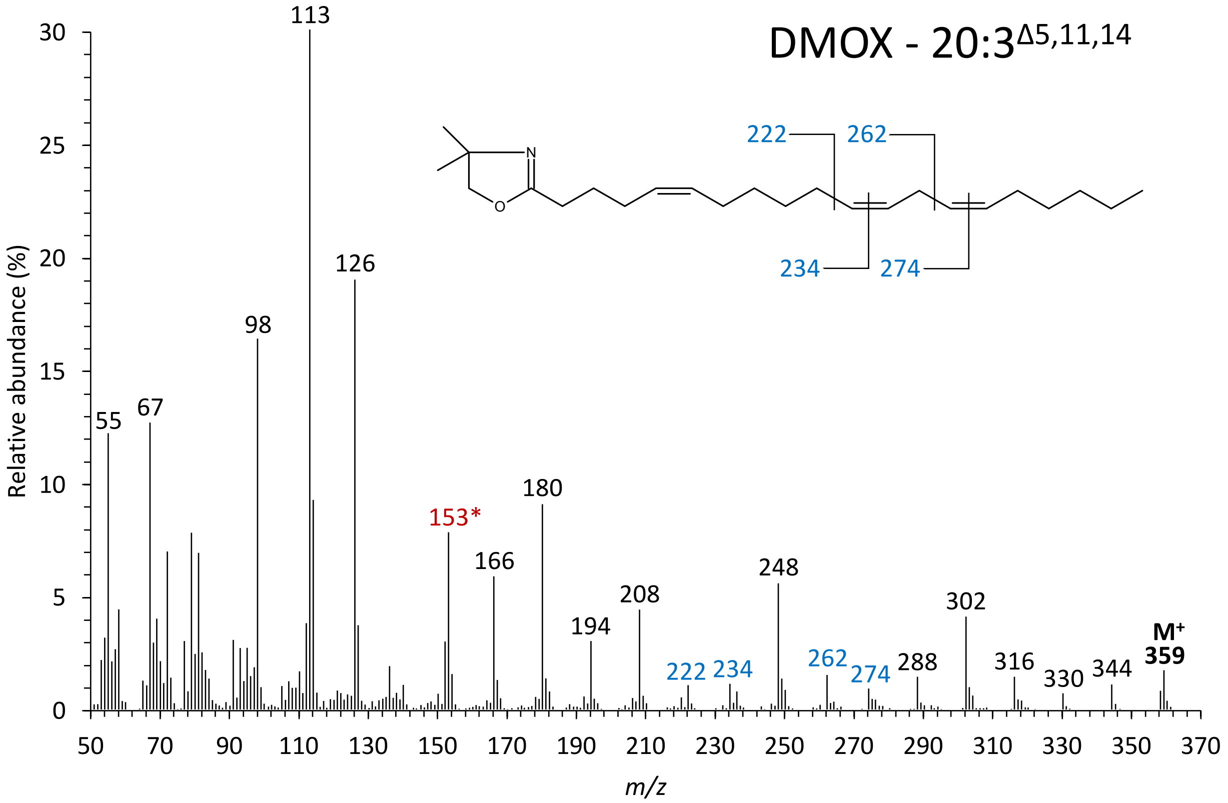


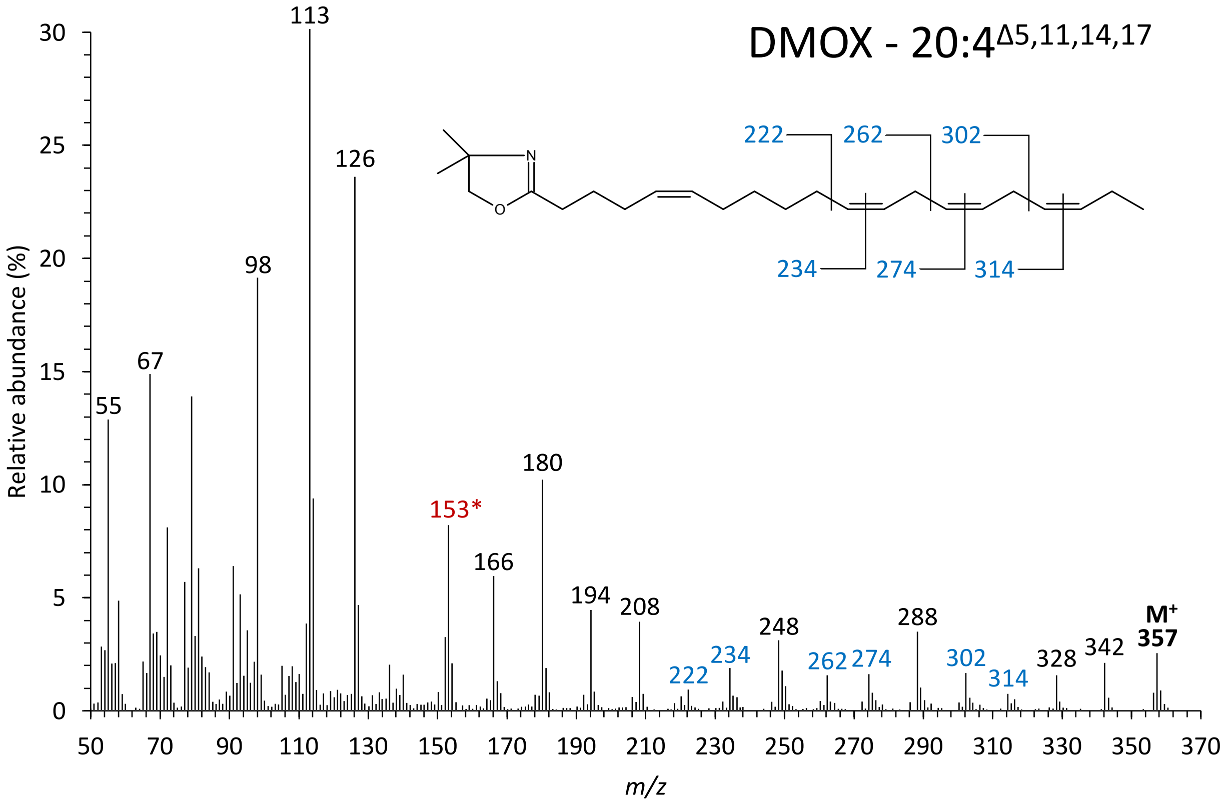
